# Mass spectrometry analysis of mouse hematopoietic stem cells and their progenitors reveals differential expression within and between proteome and transcriptome throughout adult and aged hematopoiesis

**DOI:** 10.1101/836692

**Authors:** Balyn W. Zaro, Joseph J. Noh, Victoria L. Mascetti, Janos Demeter, Benson M. George, Monika Zukowska, Gunsagar S. Gulati, Rahul Sinha, Rachel M. Morganti, Allison M. Banuelos, Allison Zhang, Peter K. Jackson, Irving L. Weissman

## Abstract

**Summary:** Hematopoietic stem cells (HSCs) are responsible for the generation of blood and immune cells throughout life. They have the unique ability to self-renew and generate more HSCs or differentiate into a progenitor cell in response to cell-intrinsic and -extrinsic stimuli. The balance of HSC fate commitment is critical for a healthy blood supply. Imbalances during hematopoiesis, which are frequent in aging, can result in hematological malignancies and pre-malignancies as well as increase risk of atherosclerosis. Given the importance of HSCs and their progenitors, they have been extensively characterized in genomic and transcriptomic studies. However, an understanding of protein expression within the HSC compartment and more broadly throughout hematopoiesis remains poorly understood, and it has been widely reported that the correlation between mRNA and proteins is more complicated than previously thought. Previous mouse mass spectrometry studies have focused either specifically on stem and the first early progenitor or broadly across mixed populations of stem and progenitor cells, which do not allow for cell-type specific protein resolution across stages of differentiation. Mass cytometry has been employed to characterize transcription factor expression in human HSCs and progenitors but does not apply an unbiased discovery approach. New mass spectrometry technology now allows for deep proteomic coverage with no more than 200 ng of sample input. We report here a proteomics resource characterizing protein expression in mouse adult and aged HSCs, multipotent progenitors and oligopotent progenitors, 12 cell types in total. We validated differential expression by flow cytometry analysis and immunofluorescence staining. Additionally, we investigated the relationship between mRNA and protein levels of individual genes in HSCs compared to progenitors through RNA sequencing studies and identified two proteins that appear to be uniquely regulated in the HSC compartment, Cpin1 and Adnp. In summary, this resource provides proteomic coverage of adult and aged hematopoietic stem cells and their progenitors and reveals changes in protein abundance between cell types, with potential future implications in understanding mechanisms for stem-cell maintenance, niche interactions and fate determination.

## Introduction

Hematopoietic stem cells (HSCs) allow for persistent renewal of blood and immune cells throughout a lifetime. They have the ability to not only self-renew but also differentiate into effector cells in response to physiological demands such as infection or bleeding (Figure 1A) (Seita and Weissman, 2010). Due to their unique capabilities and therapeutic promise in regenerative medicine, organ transplantation and hematological malignancies and pre-malignancies, the biology behind HSCs and their progeny has been of extreme interest to the scien ific and medical communities since Till and McCulloch’s first indication of their existence in 1961-68 (Weissman and Shizuru, 2008). It has since been demonstrated that HSCs can give rise to non-committed multipotent progenitor cells (MPPs) that move through differentiation, eventually committing to a more-defined cell fate in the oligopotent progenitor (OPP) compartment (Figure 1A) (Akashi et al., 2000; Kondo et al., 1997). The OPP compartment consists of fate-restricted cell types: the common myeloid progenitor (CMP), the granulocyte-macrophage progenitor (GMP), the megakaryocyte-erythroid progenitor (MEP) and the common lymphoid progenitor (CLP) (Figure 1A). Importantly, mouse and human hematopoietic cell surfaces are dramatically reengineered during differentiation. Over the past 30 years, surface proteins have been identified and differential expression phenotypically characterized, allowing for purification of phenotypically-distinct cell populations through single-cell sorting in conjunction with a panel of fluorescent antibodies (Ema et al., 2006; Kiel et al., 2005; Majeti et al., 2007; Miller and Eaves, 1997; Morrison and Weissman, 1994; Spangrude et al., 1988).

**Figure 1.**
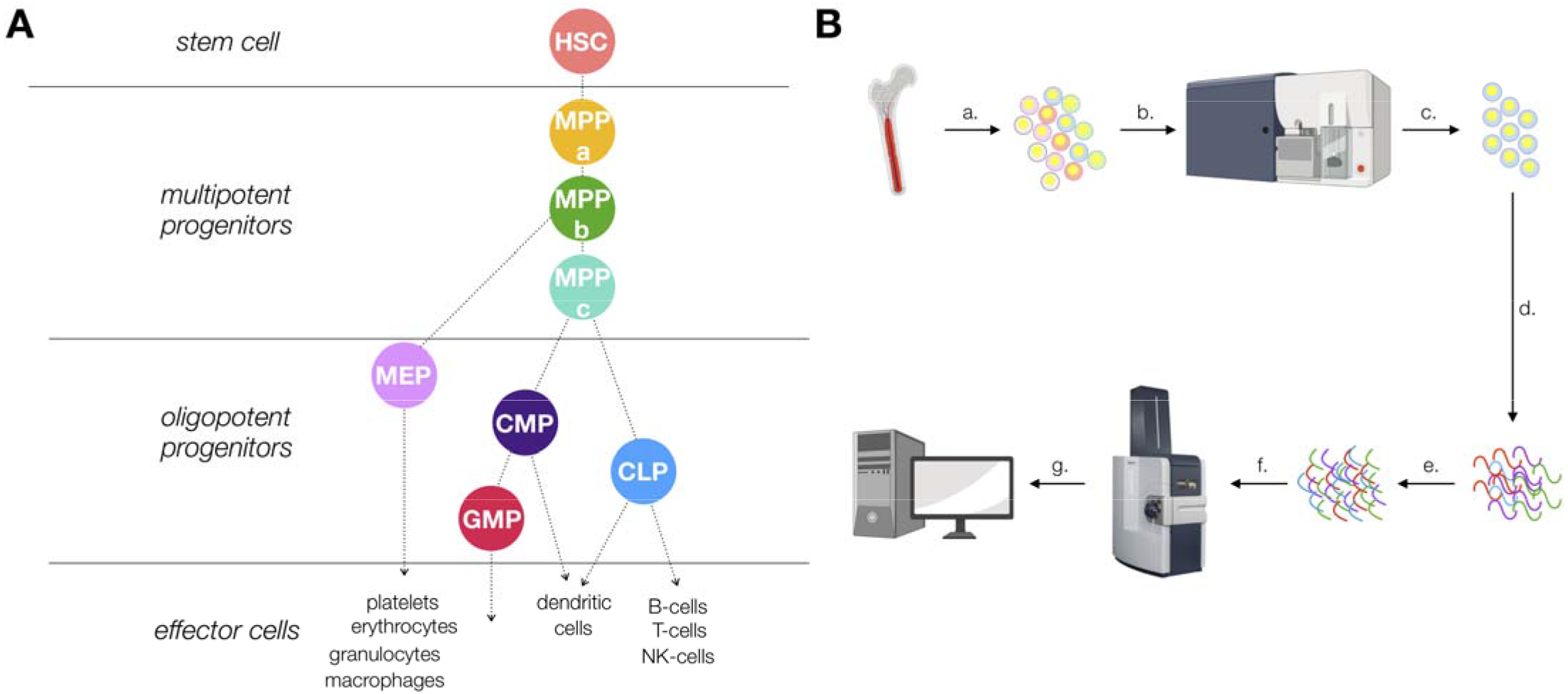
Proteomics of mouse hematopoiesis workflow. A. The hematopoietic tree. Hematopoietic stem cells (HSCs) gi e rise to multipotent progenitors (MPPs). Fate commitment arises in the oligopotent progenitor (OPP) com artment: megakaryocyte/erythrocyte progenitors (MEPs), common myeloid progenitors (CMPs), common lymphoid progenitors (CLPs) and granulocyte/macrophage progenitors (GMPs). B. Proteomic sample preparation workflow: a. bone marrow cells are isolated and a single cell suspension generated b. cells are stained with a panel of antibodies c. cells are sorted by FACS. d. cells are lysed and protein amounts normalized e. lysate is digested and desalted f. peptides are subjected to mass spectrometry analysis g. raw data is processed and analyzed to generate datasets.

Blood malignancies and cytopenias can be successfully treated with purified HSC transplants and are also used during organ transplantation to promote engraftment and tolerance (Gandy et al., 1999; Shizuru et al., 1996; Uchida et al., 1998). Transplanted purified HSCs can completely reconstitute blood lineages in conditioned mice without causing graft vs. host disease in an allogeneic setting. Harsh conditioning regimens have previously been required prior to allogeneic transplant. However, we and others have reported the use of an antibody depletion cocktail which can be used in lieu of traditional chemotherapy and irradiation strategies, increasing the potential application of hematopoietic cell transplantations well beyond otherwise-fatal malignancies (Chhabra et al., 2016; Czechowicz et al., 2007; George et al., 2019). These transformative developments further emphasize our need to completely understand HSC maintenance and fate determination and how hematopoiesis is altered during disease and aging.

The dynamics of HSC biology have revealed unique behaviors employed by these cells to maintain stem-like properties. During aging, HSC populations become less self-renewing and exhibit a myeloid-biased fate, prompting expansion of the HSC pool and an increase in CMP production (Beerman et al., 2010; Muller-Sieburg et al., 2004; 2002; Pang et al., 2011). Aged HSCs can give rise to imbalanced output and misregulated progenitor cells, resulting in the presentation of cytopenias before onset of a treatable disease (Jaiswal et al., 2014). Clonal hematopoiesis of indeterminate potential (CHIP) is associated with blood cancers, atherosclerosis and premature death(Jaiswal et al., 2014; 2017). In pre-malignancies, diseased HSCs outcompete healthy HSCs for their coveted bone marrow niche and thus become the primary drivers of blood formation (Corces et al., 2018; Jan et al., 2012; Pang et al., 2013). Investigation into the mechanisms behind such biological phenomena can provide insight into genes and proteins critical for HSC fate and function.

Given the historical challenge of collecting enough of these cells for experimental studies, much of what is known about HSCs and their progeny has been discovered through DNA microarray, bulk and single-cell RNA-sequencing and ATAC-sequencing experiments. Functional transplant studies and/or phenotypic genetic knockout mouse models have been the cornerstones of our understanding for how HSCs maintain stemness and determine their fate (Lu et al., 2011). While there is currently an abundance of genomic and transcriptomic sequencing available, very little proteomic or metabolomic investigation has been conducted (Buenrostro et al., 2018; Cabezas-Wallscheid et al., 2014; Galeev et al., 2016; Seita et al., 2012). Recent publications suggest that the hematopoietic hierarchy is more nuanced and less single-path driven than originally thought (Rodriguez-Fraticelli et al., 2018; Sanjuan-Pla et al., 2013; Yamamoto et al., 2018a; 2018b). These discoveries emphasize that there is still much to be uncovered in our understanding of HSC and progenitor biology.

Large scale mRNA and genomic sequencing studies provide insight into the consequences of perturbations in gene expression during differentiation, development, disease and aging, but we still lack a complete understanding of how genes drive function through protein expression. It has been well documented that mRNA abundance and protein expression are not well correlated (Gygi et al., 1999; Koussounadis et al., 2015; Liu et al., 2016). Additionally protein translation studies in HSCs suggest multiple modes in the regulation of protein expression, and therefore determining mRNA levels of proteins of interest is insufficient for a complete understanding of the hematopoietic protein profile in stem and progenitor compartments(Signer et al., 2014).

There are many lines of evidence that suggest HSCs exhibit minimal regulation at the genomic and transcript level. Previous reports have demonstrated that protein synthesis is tightly regulated in the HSC compartment compared to downstream progenitors and altering this rate of translation impairs HSC function (Signer et al., 2014). Rates of protein synthesis are significantly lower in HSCs compared to downstream progenitors and, importantly, these discrepancies cannot be entirely explained by differences in ribosomal RNA and total cellular RNA content (Buszczak et al., 2014; Signer et al., 2014). It has also been demonstrated that HSCs present more open chromatin compared to that of progenitors (Miyamoto et al., 2002; Yu et al., 2016). In single cell ATAC-seq data, chromatin accessibility landscapes were determined to be highly variable in early-stage hematopoietic cells, suggesting reduced regulation at the chromatin level and increased message diversity (Buenrostro et al., 2018). These data further support a hypothesis whereby protein translation is the vital regulatory step in HSC fate determination, and a complete understanding of the suite of proteins expressed by each cell type during hematopoiesis can provide further insight into the biological mechanisms at play.

Previously, Trumpp and co-workers reported comparative proteomic data between HSCs and early MPP1 progenitors (Lin^−^, Sca1^+^, cKIT^+^, CD34^+^, CD48^−^, CD150^+^), which provides insight exclusively into proteins critical to the earliest stage in hematopoiesis (Cabezas-Wallscheid et al., 2014). Palii et. al. have utilized targeted mass cytometry to characterize expression of specific transcription factors critical to human erythropoiesis (Palii et al., 2019). Proteomics of mixed populations of mouse HSC stem and progenitor cells (Lin^−^, Sca1^+^, cKIT^+^) in fetal and adult has been reported by Hansson and co-workers, but the imbalance in proteomic contribution between HSCs and progenitors in the multipotent and oligopotent compartments does not allow for further insight into the fate and function of each specific cell type (Jassinskaja et al., 2017). Currently, we lack an understanding of protein expression across early hematopoiesis, including HSCs, MPPs (MPPa, b and c) and OPPs (CMP, GMP, MEP, CLP). This knowledge gap hampers our ability to determine key players in biochemical processes critical to hematopoiesis, to identify cell-specific surface proteins for improved purification strategies and to discover novel therapeutic protein targets.

HSCs are very rare and often difficult to purify, presenting a formidable challenge for traditional biochemical methods of investigation. Approximately 0.01-0.001% of mononuclear cells in the bone marrow of adult humans and mice are HSCs, and therefore experiments requiring large amounts of highly-pure starting material, such as cell lysate, have not been technically achievable (Mayle et al., 2013). Recently the Mann laboratory reported the use of mass spectrometry instrumentation capable of increased sensitivity and improved proteomic coverage with very low amounts of protein(Meier et al., 2018). This technology allows for unprecedented coverage from samples under 200 ng, dramatically improving the feasibility of performing large-scale proteomics studies on rare cell types.

With this technology available to us and our awareness of the incomplete profiling of the hematopoietic proteome, we sought to complement and elaborate upon these data with a comprehensive unbiased proteomics database characterizing protein expression throughout the mouse adult hematopoietic hierarchy from HSCs through the multipotent and oligopotent progenitor compartments as well in aged HSCs and MPPs. The database has been validated through FACS and fluorescence microscopy experiments for proteins of interest. In comparing our proteomics database with bulk mRNA sequencing from HSCs, MPPas, MPPbs and MPPcs we identified a unique relationship between transcription and translation exclusive to the HSC compartment. Finally, these protein expression data are now integrated into a resource that allows for researchers to understand not only how mRNA levels change for genes of interest but also how protein expression is altered during adult and aged hematopoiesis.

## Results

### Optimization of sample preparation for low numbers of rare cells

Critical to our ability to deeply characterize protein expression was the development of a method by which to efficiently purify and process samples for mass spectrometry analysis from under 100,000 cells. To this end, we created a workflow whereby cells were purified by FACS using three sorting panels allowing for the isolation of 8 cells types: HSC (Lin^−^, cKIT^+^, Sca1^+^, CD34^−^, CD150^+^, Flt3^−^), MPPa (Lin^−^, cKIT^+^, Sca1^+^, CD34^+^, CD150^+^, Flt3^−^), MPPb (Lin^−^, cKIT^+^, Sca1^+^, CD34^+^, CD150^−^, Flt3^−^), MPPc (Lin^−^, cKIT^+^, Sca1^+^, CD34^+^, CD150^−^, Flt3^+^), CLP (Lin^−^, CD34^med/hi^, Flt3^+^, IL7Rα^+^, cKIT^lo^, Sca1^lo^), CMP (Lin^−^, cKIT^+^, Sca1^lo/−^, CD34^med/hi^, CD16/32^−/lo^), MEP (Lin^−^, cKIT^+^, Sca1^lo/−^, CD34^−^, CD16/32^−/lo^, CD150^+^) and GMP (Lin^−^, cKIT^+^, Sca1^lo/−^, CD34^hi^, CD16/32^hi^) (Figure 1 and Supplemental Figures 1A, 1B and 1C). Cells were purity sorted into FACS buffer (2% fetal bovine serum in PBS) and washed twice with PBS to remove any remaining serum prior to storage. Samples for mass spectrometry analysis were prepared with a minimum of 50,000 cells (Figure 1B). A commercially-available mass spectrometry sample preparation kit was utilized to ensure minimal sample loss and reproducibility and proved critical to our efforts (Figure 1B). For all adult cells and aged HSCs, at least three biological cohorts of mice were utilized for each cell type, and the sample was run in technical duplicate with 200 ng of loading material per replicate. Six total replicates were acquired for 9 cell types of interest, including aged HSCs, and 4 replicates were acquired for the 3 aged MPP populations. Each technical replicate was processed through Byonic software as an individual dataset.

### A database of proteins expressed by rare cell types

In order to generate a repository of proteins expressed by HSCs and their progenitors that are detectable by mass spectrometry, we divided each protein intensity by the total protein intensity per technical replicate and multiplied by 1 million for ease of analysis (Table 1). An average was taken within each cell type for global analyses (Table 2). Across all cell types we detected a total of 7917 genes encoding proteins expressed and detectable in HSCs and their progenitors (Tables 1 and 2). The adult HSC compartment had the least protein diversity, with 4030 proteins detected (Figure 2A). We observed a general trend in increased protein diversity during the differentiation process, with MPPs and OPPs expressing larger numbers of distinct proteins compared to HSCs (Figure 2A). This result differs from previous reports where HSCs have been demonstrated to present increased mRNA diversity in the stem cell compartment compared to their progeny(Ramos et al., 2006).

**Figure 2.**
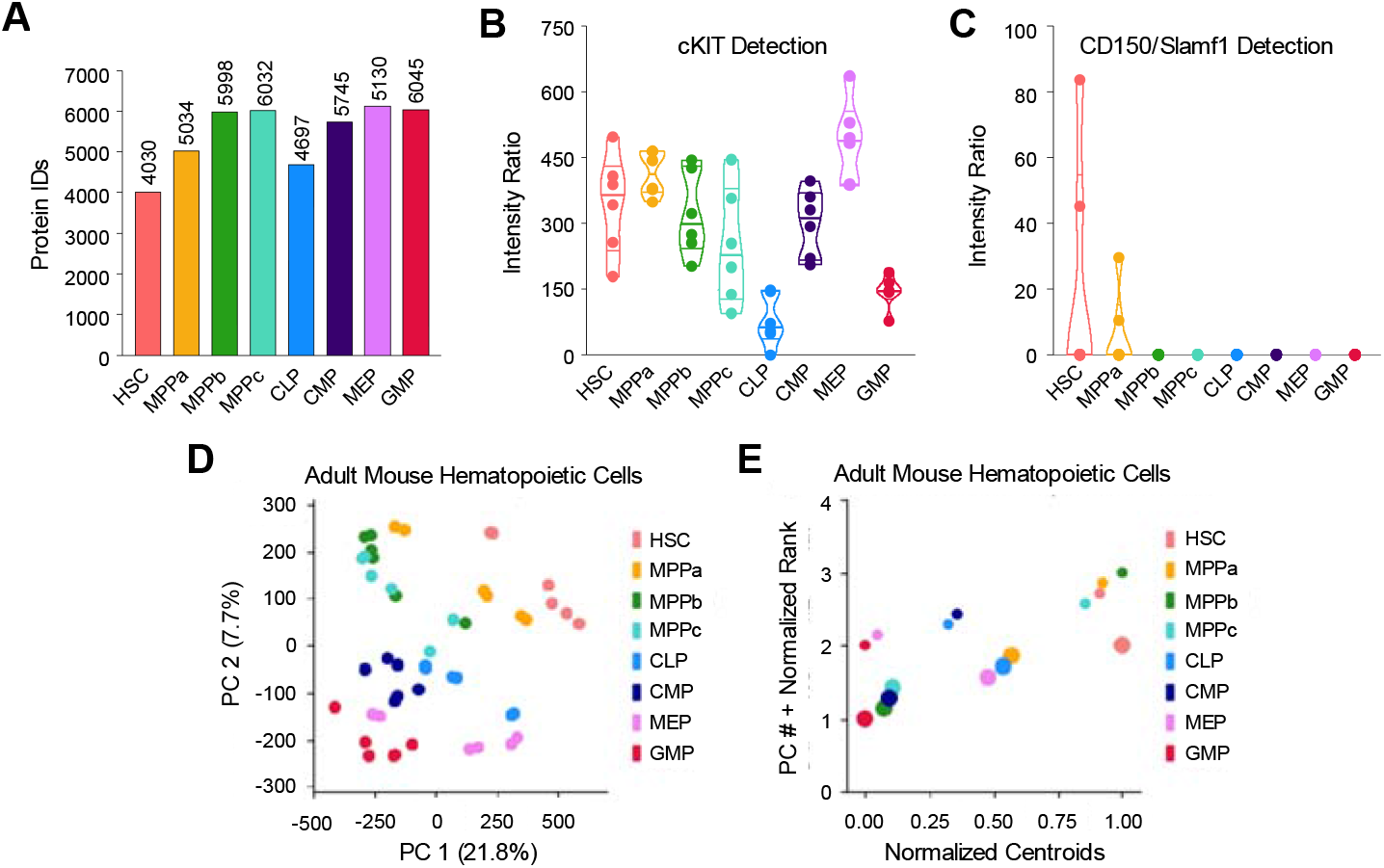
Proteomic profiling of HSCs and their progenitors. A. The number of proteins identified in each cell type across 6 replicates. B. Validation of cKIT expression in HSCs and progenitors. Violin plots show intensity ratios of cKIT expression from 6 replicates per cell type. C. Validation of CD150/Slamf1 expression exclusively in the HSC and MPPa cell types. Violin plots show intensity ratios of CD150/Slamf1 expression from 6 replicates per cell type. D. PCA Plots of all replicates per cell type display protein expression throughout the hematopoietic tree. E. 1-dimensional PC plots show which components are key drivers of segmentation between cell types and cell compartments. Centroids are normalized representatives of all 6 replicates and are scaled in size with respect to the percent variation explained by each principal component.

### Characterization of protein expression in HSCs and their progenitors

With our database generated, we next validated detection of known markers of stem and progenitor cells, including cKIT and CD150/Slamf1 (Figures 2B and 2C). As expected, cKIT was detected across all cell types with expression high in HSCs, MPPs and lower in CLPs and GMPs. CD150/Slamf1 expression was exclusive to HSC and MPPa compartments, an attestation to the purity of these sorted samples. The cell-cycle associated protein ki67 was lowly detected in the HSC compartment with a steady increase in expression in the MPP compartment and highest in the OPPs (Supplemental Figure 1D). Principal component analysis (PCA) revealed exquisite distribution of adult hematopoietic stem and progenitor cells during differentiation (Figure 2D, Table 4). Excitingly, each component revealed unique attributes driving differentiation: component 1 isolates HSCs from all other cell types, whereas component 2 separates stem and multipotent progenitors from the more-committed oligopotent progenitor compartment (Figure 2E and Table 4).

### Validation of differential expression of proteins of interest

To validate additional targets using non-mass spectrometry techniques, we performed FACS analysis and fluorescence microscopy. The endothelial surface adhesion molecule (ESAM) has previously been shown to be highly expressed by HSCs(Ishibashi et al., 2016; Ooi et al., 2009; Yokota et al., 2009). In our dataset, ESAM expression was very high in HSCs and MPPas with decreased expression in MPPbs and no detection for the remaining cell types (Figure 3A). This result was recapitulated in flow cytometry analysis of ESAM expression, further supporting the quality of the proteomics dataset. (Figure 3B and Supplemental Figure 2). We also characterized differential expression of the regulatory glycolytic enzyme phosphofructokinase (Pfkl) by fluorescence microscopy and mass spectrometry (Figures 3C and D). Pfkl expression was consistently decreased in the HSC compartment compared to MPPas and MPPbs.

**Figure 3.**
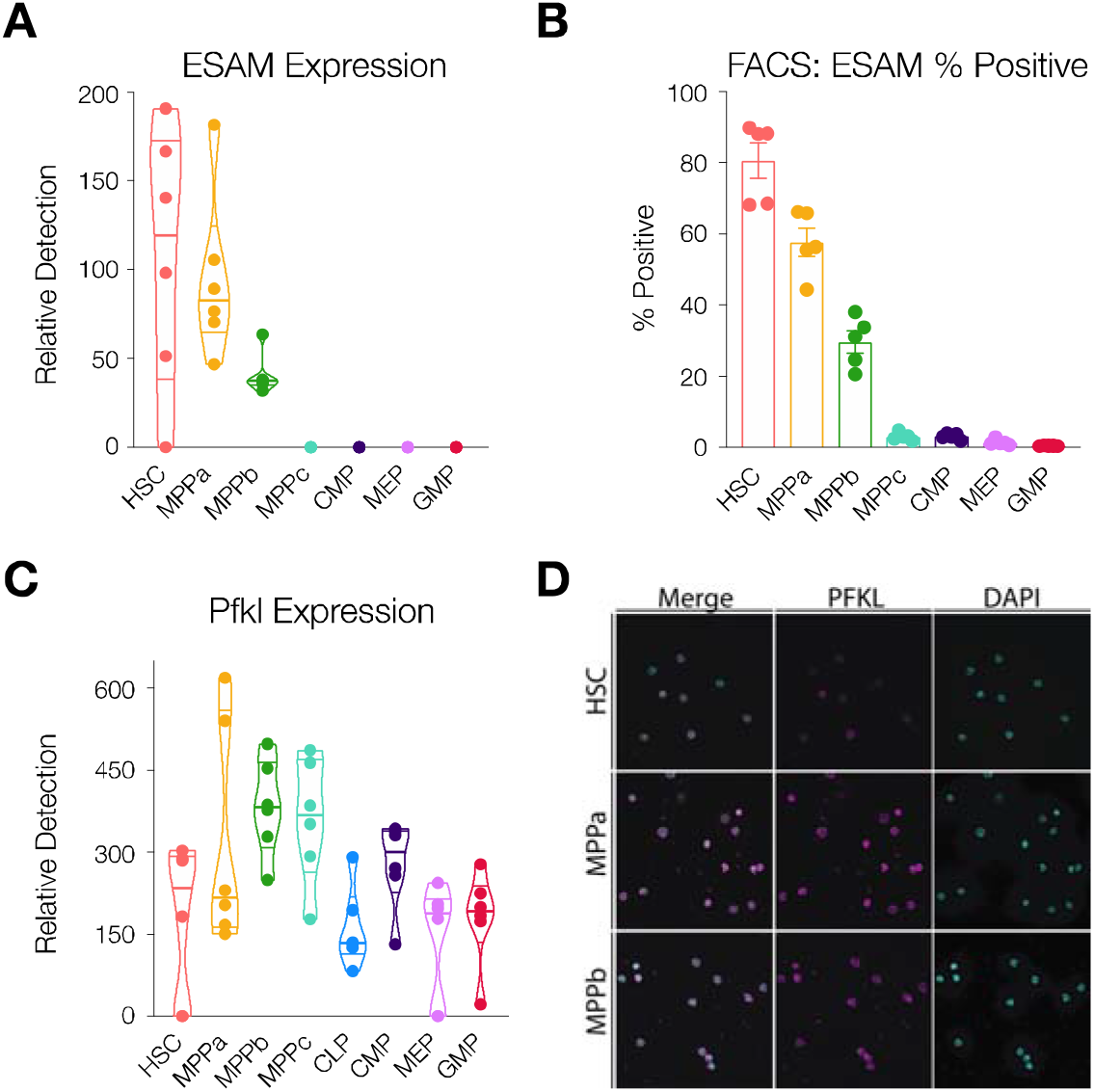
Validation of proteins of interest. A. ESAM expression in HSCs and progenitors. Violin plots show intensity ratios of ESAM expression from 6 replicates per cell type. B. ESAM expression as determined by flow cytometry analysis. N = 5 mice (3 male, 2 female). C. A. Pfkl expression in HSCs and progenitors. Violin plots show intensity ratios of Pfkl expression from 6 replicates per cell type. D. Fluorescence microscopy with antibody staining of Pfkl in HSCs, MPPas and MPPbs. N = 5 mice.

This finding prompted a deeper investigation into changes to expression of glycolytic enzymes. For enzymes critical to glycolysis, we noted that many enzymes involved in the pathway were more-highly expressed in HSCs or comparably expressed across HSCs and early progenitors with the exception of the rate limiting step, phosphofructokinase (Pfk) (Figure 3C and Supplemental Figure 3). It has been previously reported by Ito and co-workers that HSCs undergo anaerobic glycolysis in order to protect themselves from the likely hypoxic conditions of their bone marrow niche, and, while our studies cannot assess the activity of these enzymes, these proteomic expression data further support a glycolytic advantage in the HSC compartment(Ito and Suda, 2014). Takubo et al. have also reported that PDK2- and PDK4-mediated regulation of anaerobic glycolysis in long-term HSCs modulates stemness and arrests cell cycle(Takubo et al., 2013). While we identify PDK2 and PDK3 in our datasets, we did not observe increased expression of these proteins, as detectable by mass spectrometry (Tables 1 and 2). This discrepancy may be due to several explanations, including low rates of detection and differences in selection markers used for each cell type. Nevertheless, differential expression of glycolytic enzymes between HSCs and progenitors remains consistent throughout all reports.

We also compared our HSC (Lin^−^, cKIT^+^, Sca1^+^, CD34^−^, CD150^+^, Flt3^−^) and MPPa (Lin^−^, cKIT^+^, Sca1^+^, CD34^+^, CD150^+^, Flt3^−^), data to that of Trumpp and co-workers’ comparative proteomics data between HSC (Lin^−^, Sca1^+^, cKIT^+^, CD34^−^, Flt3^−^, CD48^−^, CD150^+^) and MPP1 (Lin^−^, Sca1^+^, cKIT^+^, CD34^+^, Flt3^−^, CD48^−^, CD150^+^) cell types (Table 5) (Cabezas-Wallscheid et al., 2014). Of the 49 differentially expressed proteins identified by Cabezas-Wallscheid et. al. that were also detected in our datasets, 35 were consistent across both experimental methods (at least 2-fold more frequently detected in HSC or early progenitor in our data or denoted as differentially expressed per Cabezas-Wallscheid et. al.). While most of these data were consistent across both experiments, such as HSC-enriched expression of Igf2bp2 and Hmga2, our additional cell-type coverage allows for a deeper resolution into differential expression throughout the hematopoietic tree (Supplemental Figures 4A and 4B) (Nishino et al., 2008; 2013). For example, Igf2bp2 is exclusively detected in HSCs in our datasets across all cell types, whereas Hmga2 is simply more abundant in the HSC compartment (Supplemental Figure 4B).

### Identification of further potential proteins of study

Over 40% of proteins were detected across all cell types (Figure 4A). With deep proteome coverage and analysis of progenitor populations, we were also able to identify proteins uniquely absent or uniquely detected in a single cell type (Figure 4A and Table 3). To validate that differential expression profiles were not predominantly due to global differences in protein detection across cell types, we looked at expression of the housekeeping protein Hprt1 (Figure 4B). Importantly, protein expression was consistent across all cell types characterized. In investigating uniquely expressed proteins, particularly the 340 proteins exclusively expressed in the GMP compartment, we identified GMP-specific proteins including CCAAT-enhancer-binding protein ε(Cebpe), Adhesion G Protein-Coupled Receptor G3 (Adgrg3) and Membrane-spanning 4-domains, subfamily A, member 3 (Ms4a3), all of which have previously been reported to be associated with macrophage and granulocyte-specific lineage commitment (Tables 1-3) (Goardon et al., 2011; Hsiao et al., 2018; Ishibashi et al., 2018; Nakajima et al., 2006). In the HSC compartment, Igf2bp2 was detected as a uniquely expressed protein, as has previously been identified as differentially expressed by HSCs compared to MPP1s and MPPas (Supplemental Figure 4A) (Cabezas-Wallscheid et al., 2014). We also identified 619 proteins that were detected in all other cell types besides HSCs (Figure 4A). Anamorsin (Cpin1), a protein encoded by the gene Ciapin1, and the transcription factor Activity-dependent neuroprotector homeobox (Adnp) were identified throughout MPP and OPP compartments but notably absent in HSCs (Figure 4C). Cpin1 is a protective anti-apoptotic protein that is required for maturation of erythroid cells but has been shown to not affect the number of stem and early progenitors cells(Shibayama et al., 2004). Adnp is a transcriptional regulator implicated in neural development that also affects erythropoiesis(Dresner et al., 2012; Mandel et al., 2007).

**Figure 4.**
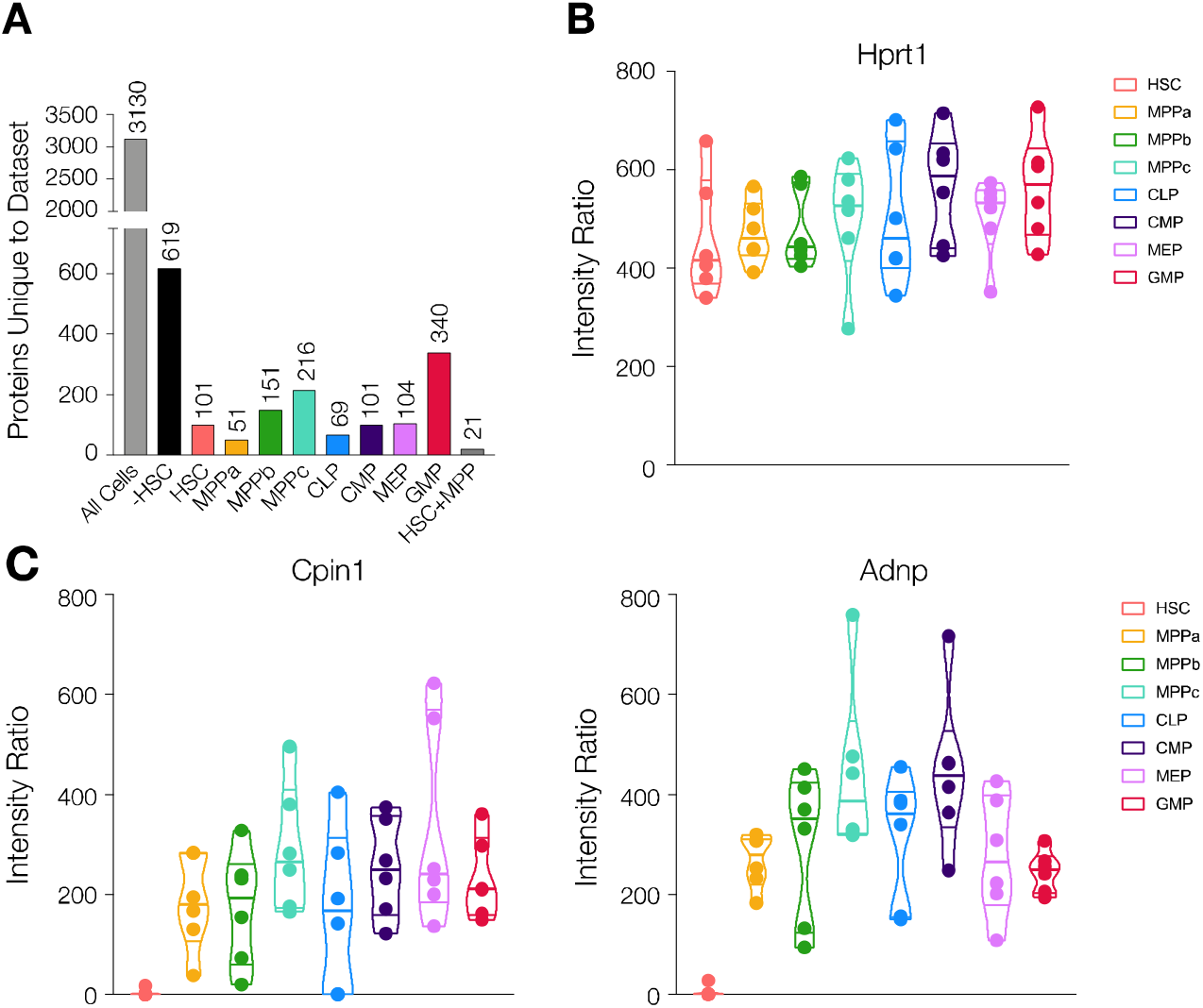
Investigation of proteins uniquely present or absent in each cell type. A. Number of proteins uniquely detected in each indicated cell type. - HSC denotes all cell types besides HSCs. HSC+MPP denotes all HSCs and MPPs (a-c). B. Violin plots for intensity ratios of the housekeeping protein Hprt1 across 6 replicates per cell type. C. Violin plots for intensity ratios of Cpin1 and Adnp across 6 replicates per cell type.

### Characterization of protein expression in Aged HSCs and MPPs

Blood formation during aging is marked by a myeloid bias and higher frequency but lower engraftment per HSC transplanted as described previously(Jaiswal et al., 2014; Morrison et al., 1996; Pang et al., 2011). However, to the best of our knowledge, no mass spectrometry-based proteomics experiments have characterized protein expression changes in the HSC compartment during aging. Using our sort schemes and sample preparation methods, we purified and processed HSCs and MPPs from mice no less than 24 months of age (Figures 1 and Supplemental Figures 1A, 1B and 1C). Data analysis revealed expression of 5434 proteins in Aged HSCs (Figure 5A, and Tables 1 and 2). PCA analysis demonstrated the high similarity between adult and aged HSCs as compared to progenitor cells but also important differences across both component 1, where aged HSCs lose the distinctness of adult HSCs, and component 2, where aged HSCs seem to occupy a unique protein signature in comparison to both adult stem and progenitor compartments (Figure 5B and Table 6). As expected, cKIT was consistently detected in all 4 cell types while CD150/Slamf1 was found in HSCs and MPPas exclusively (Figure 4C). Earlier FACS separation of CD150^hi^ and CD150^lo^ HSCs revealed CD150^hi^ HSCs are myeloid biased, and this sub-population increases most-dramatically in aged mice (Beerman et al., 2010). Our proteomics data revealed similar variations in CD150 expression levels, with the range lowest in adult HSCs. Additionally, cKIT expression was much higher in the aged cells compared to their adult counterparts, which has also been observed by others (Beerman et al., 2010; Mann et al., 2018). Finally, Ki67 expression was still lower in the aged HSC compartment compared to that of adult and aged downstream progenitors (Supplemental Figure 5A).

**Figure 5.**
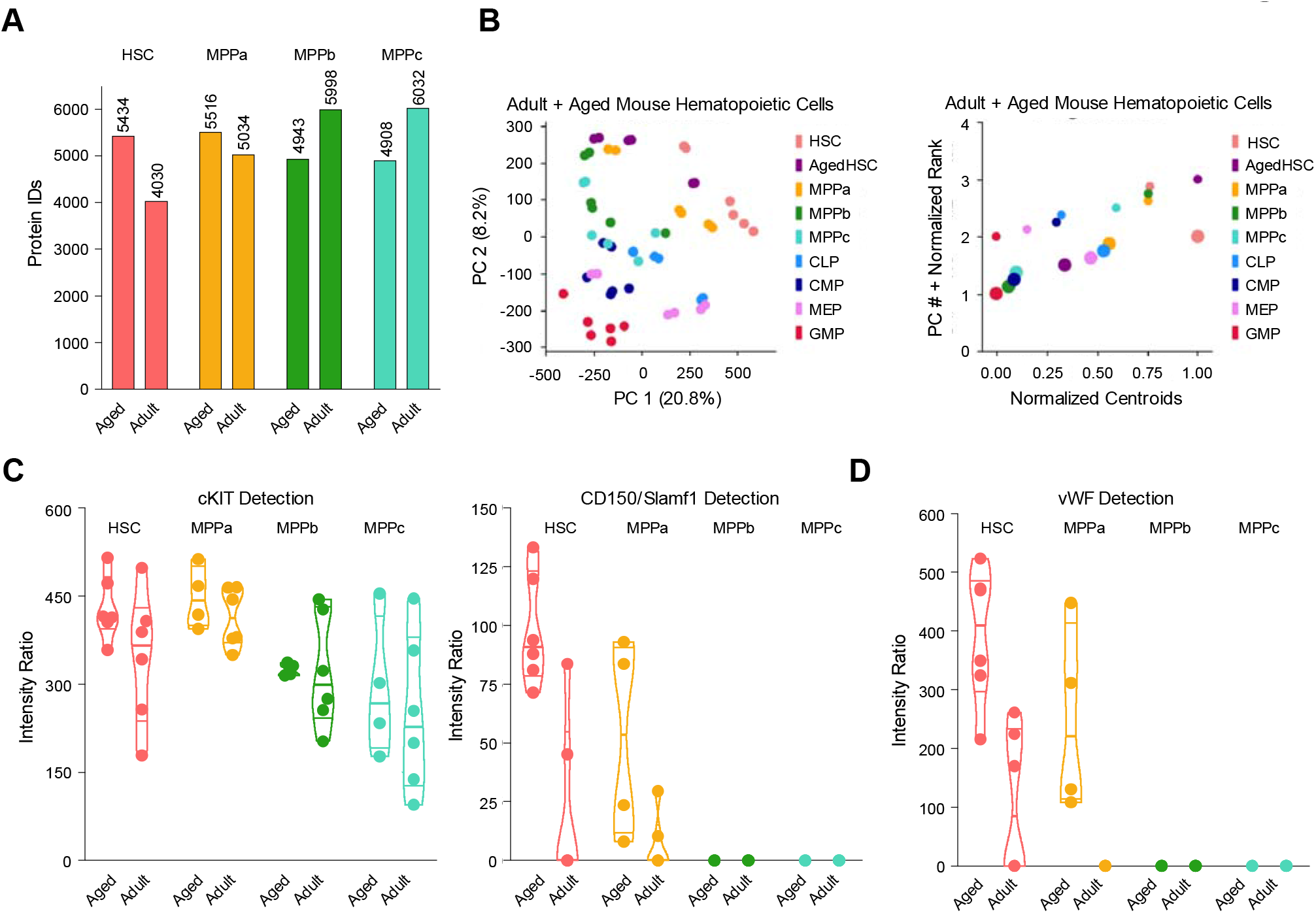
Proteomics of aged HSCs and MPPs. A. Proteins detected by mass spectrometry for the indicated aged and adult cell type. For all adult cells and aged HSCs, N = 6. For Aged MPPs, N = 4. B. PCA plots of adult HSCs and progenitors and aged HSCs. N = 6. 1-dimensional PC plots show which components are key drivers of segmentation between cell types and cell compartments. Centroids are normalized representatives of all replicates and are scaled in size with respect to the percent variation explained by each principal component. C. Validation of cKIT and CD150/Slamf1 expression in adult and aged HSCs and MPPs. Violin plots show intensity ratios of cKIT and CD150/Slamf1 expression from 6 replicates per cell type (adult cells and aged HSCs) and 4 replicates per cell type (aged MPPs). D. Validation of von-Willebrand factor (vWF) expression in adult and aged HSCs and MPPs.

Given the abundance of functional studies that have been conducted to identify genes implicated in HSC fate determination and stemness, we were interested to compare our findings of differentially expressed proteins during adult vs. aged hematopoiesis with previous reports. von-Willebrand Factor (vWF) expression is associated with myeloid and platelet biases during hematopoiesis, and we also detected increased expression of vWF in aged HSCs and MPPas (Figure 5D) (Grover et al., 2016; Mann et al., 2018; Pinho et al., 2018; Sanjuan-Pla et al., 2013). Integrin surface proteins are critical in mediating a pro-inflammatory response that can elicit bias in HSC fate determination, and such proteins are also well documented to exhibit increased expression during aging in addition to inflammatory events (Gekas and Graf, 2013; Haas et al., 2015; Mann et al., 2018; Pang et al., 2011). We detected ITGA2b (CD41) expression in aged HSCs, which has been demonstrated to induce a myeloid bias, but, similar to Mann et. al., we did not see a significant difference between aged and adult (Tables 1 and 2) (Gekas and Graf, 2013; Mann et al., 2018). However, expression of the complementary signaling molecule ITGB3 (CD61) increased significantly in the aged compartments of HSCs, MPPas and MPPbs, as described previously by Regev, Baltimore and co-workers (Supplemental Figure 5B) (Mann et al., 2018). Finally, we investigated how protein expression of Cpin1 and Adnp changes during aging and determined that expression of both proteins is rescued in the aged HSC compartment (Supplemental Figure 5C).

### mRNA abundance comparison

Globally, mRNA expression and protein abundance are not well correlated across yeast and higher eukaryotes(Liu et al., 2016). We were interested to determine if there were changes in the relationship between mRNA and protein during hematopoiesis, both broadly across the proteome as well as for specific proteins of interest. Bulk mRNA sequencing was conducted from adult mouse HSCs, MPPas, MPPbs and MPPcs as described previously (Table 7) (Moraga et al., 2015). Spearman correlation coefficients (ρ) were calculated to determine the degree of monotonic relationship between mRNA and protein for HSCs, MPPas, MPPbs and MPPcs (Figure 6A). Correlation was lowest for the HSC compartment, which supports data generated by Signer, Morrison and co-workers suggesting that protein translation is uniquely regulated in the HSC compartment at least in part through a mechanism other than altered gene transcription (Signer et al., 2014). We next compared how the protein vs. mRNA relationship is altered across HSCs and MPPs for genes expressed by all 4 cell types. Again, we detected a difference in the expression ratios, where HSCs present reduced protein translated per message available compared to the MPP compartment (Figure 6B). Finally, we investigated if the relationship of mRNA and protein was altered across cell types for proteins of interest Cpin1 and Adnp. While mRNA transcripts of Cpin1 and Adnp were detected in the HSC compartment and at comparable levels to that of MPPs, protein expression was markedly reduced (Figure 6C and Supplemental Figure 6). Importantly, the mRNA and protein expression levels of our housekeeping protein Hprt1 was consistent across all cell types.

**Figure 6.**
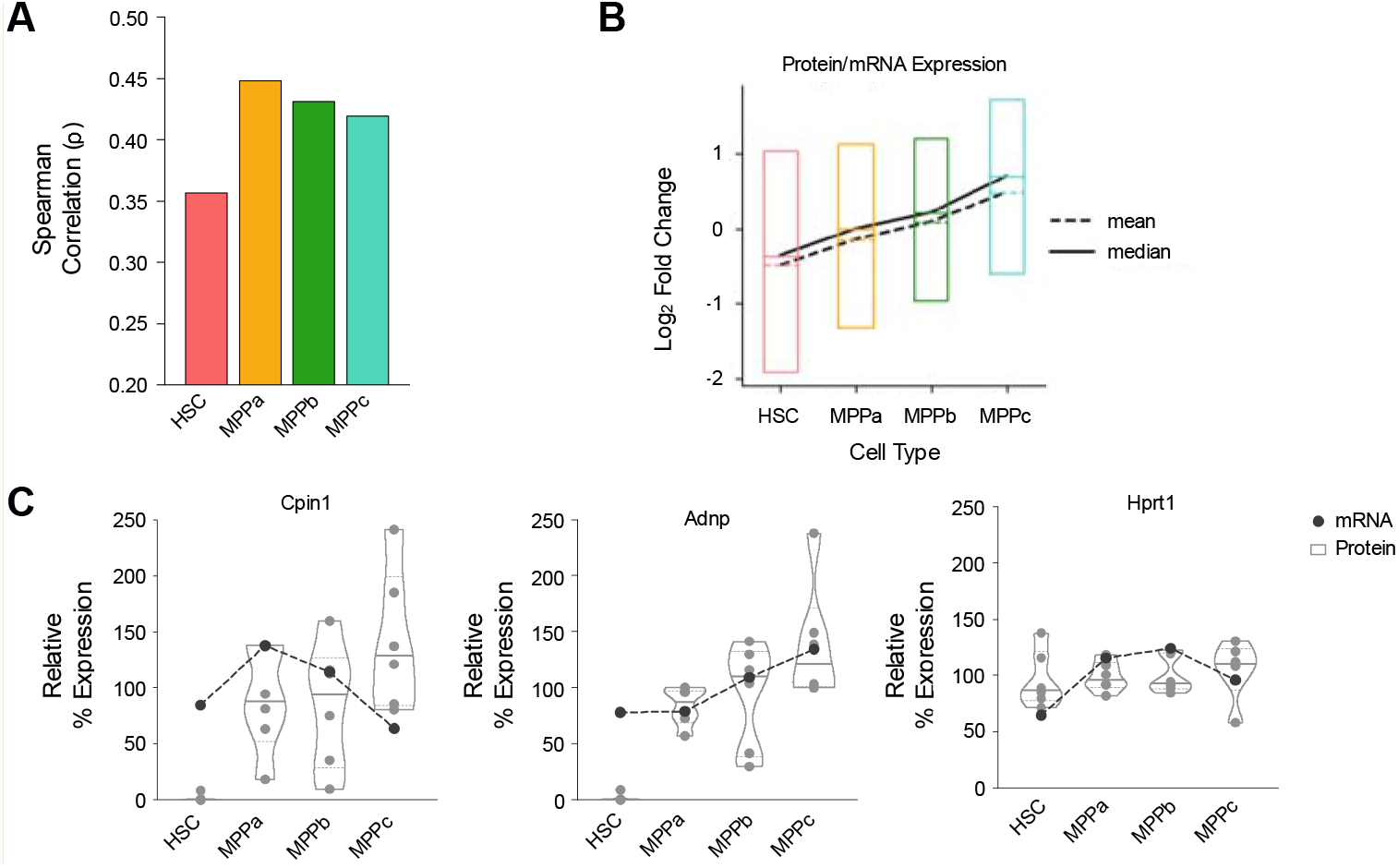
mRNA and protein have a unique relationship in the HSC compartment. A. Spearman correlation for each cell type as calculated by protein intensity ratios and mRNA levels. B. Log_2_-fold change in Protein vs. mRNA values for each cell type for proteins detected in all 4 cell types. C. Relative expression profiles of mRNA and protein for proteins/genes of interest (Cpin1/Ciapin1, Andp) and the housekeeping protein Hprt1.

## Discussion

This resource provides unprecedented coverage of protein expression for adult and aged hematopoietic stem cells and their progenitors. We validated the quality of our data via multiple methods. Our data are consistent with the expression of established surface markers, such as cKit and CD150/Slamf1, that have been validated to functionally separate HSCs and progenitors. PCA visualizes the clustering of cell types such that it is biologically meaningful for each principal component. We also confirm the validity of the total intensity readouts via FACS analysis and microscopy of select genes including ESAM and Pfkl. Many of the protein expression profiles identified validate functional and qualitative studies reported by previous groups, for example ESAM, glycolytic enzymes, Igf2bp2, and Hmga2. The increased expression of CD150/Slamf1 and vWF in the aged HSC and MPPa compartment is consistent with previous observations of increased myeloid bias in aged HSCs. The fewer protein per mRNA in HSCs corroborates decreased rates of protein synthesis in the HSC compartment compared to downstream progenitors and may explain a higher RNA complexity compared to that of protein. Put simply, some RNAs may not be translated, and their presence could reflect the opening of their chromatin rather than the need for these proteins to be translated within the HSC compartment. Our PCA also supports the idea that HSCs exhibit minimal regulation at the genomic and transcript level. The bottom 250 proteins contributing to component 1 of the adult PCA, that is proteins less expressed in HSCs compared to all other cell types, enriched for proteins involved in “chromatin silencing”, “histone modification”, and “mRNA processing”. Taken together with previous literature reports on protein translation and chromatin structure in the HSC compartment, these data suggest a new hypothesis in the regulation of stem maintenance and HSC homeostasis (Figure 7). RNA levels (mRNA and ribosomal) are comparable in HSCs compared to progenitor cells, and it has been reported that HSCs have more open chromatin than MPPs, suggesting an increased plasticity in gene transcription (Buenrostro et al., 2018). Conversely, protein translation is markedly reduced and highly sensitive to perturbations in the HSC compartment, and we now report a lower correlation between mRNA and protein (Signer et al., 2014). We therefore propose that perhaps primary regulatory mechanisms of stemness shift downstream of gene transcription towards protein translation specifically in the HSC compartment (Figure 7).

**Figure 7.**
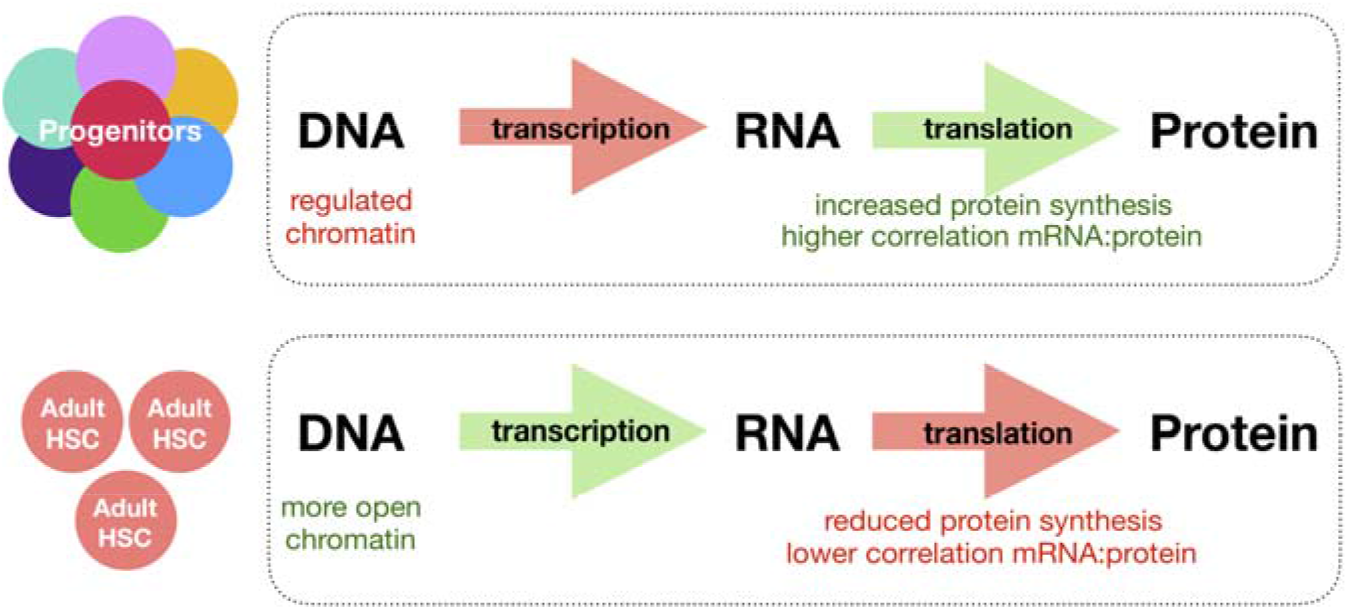
Working hypothesis for translational control in adult HSCs. HSCs exhibit more open chromatin and more diversity of mRNA. Protein translation is attenuated in HSCs resulting in a lower correlation between mRNA and protein.

While these data provide a deeper understanding of proteins expressed during adult and aged hematopoiesis that are currently detectable by mass spectrometry, we caution that this resource is by no means a complete list of proteins expressed throughout early hematopoiesis. As mass spectrometry methods continue to allow for improved data coverage with low amounts of protein, deeper characterization will become possible. It also will open up the opportunity to further segment stem and progenitor cell fractions, such as the fractionation of the HSC compartment based on the different stages of cell cycle. This can be important for our analysis of protein expression changes throughout early hematopoiesis, as HSCs are more quiescent than downstream progenitors, which can contribute to differential expression of proteins as a consequence of cell cycle. However, Signer, Morrison and co-workers have reported that differences in the rates of protein translation between HSCs and MPPs cannot be entirely explained by cell-cycle. Our data reveal global differential regulation in protein expression, some of which will be cell cycle-dependent as well as -independent (Signer et al., 2014). Given the dearth of proteomic information currently available for these rare cells, this resource reveals previously-uncharacterized suites of proteins detectable in adult and aged hematopoietic stem cell and progenitors. The nature of these expansive data allows not only for the identification of novel surface markers of each cell type but also provides an understanding of intracellular regulatory proteins of gene transcription, protein translation and metabolism that contribute to stem- and progenitor-cell survival, fate commitment and function.

## Acknowledgements

We thank Linda Quinn, Aaron McCarty, and Teja Naik for technical assistance. We thank Catherine Carswell-Crumpton, Patty Lovelace and Stephen Weber for flow cytometry assistance. This study was supported by the California Institute for Regenerative Medicine RT3-07683, the Ludwig Cancer Foundation and NIH/NCI Outstanding Investigator Award R35CA220434 (to I.L.W.); the Program in Translational and Experimental Hematology T32 from the National Heart, Lung, and Blood Institute T32HL120824 and the American Cancer Society Fellowship PF-15-142-01-CDD (to B.W.Z.); and PHS grant CA09302, awarded by the National Cancer Institute, DHHS (to B.M.G.).

## Author Contributions

B.W.Z. conceived of the project, designed experiments, performed experiments and analysis and wrote the manuscript. J.J.N. performed experiments and analysis and wrote the manuscript. V.L.M. performed experiments and analyzed the data. J.D. analyzed raw mass spectrometry data. B.M.G. and M.Z. designed and performed experiments. G.S.G. analyzed data. R.S. and R.M.M. performed and analyzed bulk RNA-sequencing data. A.M.B. and A.Z. processed mice. P.K.J. provided mass spectrometry resources. I.L.W. oversaw experiments and wrote the manuscript.

