## Supplementary material for "Mass spectrometry analysis of mouse hematopoietic stem cells and their progenitors reveals differential expression within and between proteome and transcriptome throughout adult and aged hematopoiesis": Methods

**STAR Methods**

**LEAD CONTACT AND MATERIALS AVAILABILITY**

Further information and requests for resources should be directed to and will be fulfilled by the Lead Contact, Irving Weissman. This study did not generate new reagents.

**EXPERIMENTAL MODEL AND SUBJECT DETAILS**

*Animals*

An in-house C57BL/6 strain of mice was used for collection of bone marrow-derived adult HSCs and progenitors at 8-14 weeks of age. For aged mouse studies, C57BL/6 mice (24-27 months) were a gift from Charles Chan. An equal number of male and female mice were used across all experiments. For adult studies, 50 mice were used for each biological cohort. For aged studies, 2 mice were used for each biological cohort. Mice for all experiments were immunocompetent and group housed in an AAALAC certified barrier facility. The light cycle in the facility is 12h on/12h off. All experiments were performed according to guidelines established by the Stanford University Administrative Panel on Laboratory Animal Care.

**METHOD DETAILS**

*Isolation of mouse bone marrow cells*

Mice were euthanized and hips, femurs, tibia, humeri and spine harvested. Bones were cleaned and crushed with a mortar and pestle to retrieve resident bone marrow cells with FACS buffer (2% fetal bovine serum (FBS) in phosphate buffered saline (PBS, pH 7.4, calcium and magnesium free). Cells in FACS buffer were passed through at 40 μ filter and pelleted (1,300 rpm x 5 min, 4°C).

*Isolation and purification of mouse HSCs and MPPs*

Bone marrow cell pellets were resuspended in 800 μL ice-cold FACS buffer per mouse. Miltenyi cKIT enrichment beads were added (15 μL per mouse) and incubated for 15 min at 4°C. Upon incubation completion cells were washed with 10 mL ice-cold FACS buffer per mouse and pelleted (1,300 rpm x 5 min, 4°C. Cell samples were resuspended in 1 mL ice-cold FACS buffer per mouse and loaded on to a Miltenyi MACS magnetic separation column and the column washed 3 x 3 mL FACS buffer. cKIT-positive cells were eluted according to manufacturer’s protocol and pelleted (1,300 rpm x 5 min, 4°C). Enriched cells were resuspended in FACS buffer (100 μL per mouse) and Anti-CD34 FITC (clone RAM34; 5.0 μg/mL final concentration) was added. Cells were incubated for 45 min on ice prior to addition of the remaining antibodies: Anti-Lineage Cocktail A700 (5 μL per mouse), Anti-cKIT APC-eFluor780 (clone 2B8; 2 μg/mL final concentration), Anti-Sca1 PE-Cy7 (clone D7; 2 μg/mL final concentration), Anti-CD150 APC (clone TC15-12F12.2; 2 μg/mL final concentration), and Anti-Flt3 PerCP-eFluor710 (clone A2F10; 2 μg/mL final concentration). The cells were incubated on ice for an additional 30 min with the complete cocktail prior to washing with FACS buffer to remove excess antibody. Cell were pelleted (1,300 rpm x 5 min, 4°C) and resuspended in fresh FACS buffer containing SYTOX Blue (1 mL, 1:3000) prior to FACS. Samples were sorted on a BD FACS Aria.

*Isolation and purification of mouse CMP, GMP, MEPs*

Filtered and pelleted marrow cells were resuspended in ACK lysis buffer (1 mL) and allowed to incubate for 5 min at ambient temperature. Lysis was then quenched with 10 mL of FACS buffer and cells pelleted (1,300 rpm x 5 min, 4°C). RBC-depleted bone marrow cell pellets were resuspended in 800 μL ice-cold FACS buffer per mouse. Miltenyi Lineage depletion beads were added (100 μL per mouse) and incubated for 10 min at 4°C. Upon incubation completion cells were loaded on to a Miltenyi MACS magnetic separation column and the column washed 3 x 3 mL FACS buffer. Flow-through was pelleted (1,300 rpm x 5 min, 4°C). Depleted cells were resuspended in FACS buffer (100 μL per mouse) and Anti-CD34 FITC (clone RAM34; 5.0 μg/mL final concentration) was added. Cells were incubated for 45 min on ice prior to addition of the remaining antibodies: Anti-Lineage Cocktail A700 (3 μL per mouse), Anti-cKIT APC-eFluor780 (clone 2B8; 2 μg/mL final concentration), Anti-Sca1 PE-Cy7 (clone D7; 2 μg/mL final concentration), Anti-CD150 APC (clone TC15-12F12.2; 2 μg/mL final concentration), and Anti-CD16/32 BV395 (2.4G2; 2 μg/mL final concentration). The cells were incubated for an additional 30 min on ice with the complete cocktail prior to washing with FACS buffer to remove excess antibody. Cell were pelleted (1,300 rpm x 5 min, 4°C) and resuspended in fresh FACS buffer containing SYTOX Blue (1 mL, 1:3000) prior to FACS. Samples were sorted on a BD FACS Aria.

*Isolation and purification of mouse CLPs*

Filtered and pelleted marrow cells were resuspended in ACK lysis buffer (1 mL) and allowed to incubate for 5 min at ambient temperature. Lysis was then quenched with 10 mL of FACS buffer and cells pelleted (1,300 rpm x 5 min, 4°C). RBC-depleted bone marrow cell pellets were resuspended in 800 μL ice-cold FACS buffer per mouse. Miltenyi Lineage depletion beads were added (100 μL per mouse) and incubated for 10 min at 4°C. Upon incubation completion cells were loaded on to a Miltenyi MACS magnetic separation column and the column washed 3 x 3 mL FACS buffer. Flow-through was pelleted (1,300 rpm x 5 min, 4°C). Depleted cells were resuspended in FACS buffer (100 μL per mouse). Cells were incubated on ice for 30 min with the following antibodies: Anti-Lineage Cocktail A700 (3 μL per mouse), Anti-cKIT APC-eFluor780 (clone 2B8; 2 μg/mL final concentration), Anti-Sca1 PE-Cy7 (clone D7; 2 μg/mL final concentration), Anti-Flt3 PerCP-eFluor710 (clone A2F10; 2 μg/mL final concentration), and Anti-IL7Ra APC (clone A7R34; 2 μg/mL final concentration). The cells were incubated on ice for an additional 30 min with the complete cocktail prior to washing with FACS buffer to remove excess antibody. Cell were pelleted (1,300 rpm x 5 min, 4°C) and resuspended in fresh FACS buffer containing SYTOX Blue (1 mL, 1:3000) prior to FACS. Samples were sorted on a BD FACS Aria.

*Isolation and purification of human stem and progenitor cells*

Fresh mononuclear cells were resuspended in to appropriate volume of FACS buffer as indicated in the Miltenyi Biosciences CD34 microbead enrichment kit. Manufacturer’s protocols using LS columns and magnetic separation were followed for the CD34 enrichment with no further modifications. Enriched cells were pelleted and resuspended in 100 μL per sample (up to 10 million CD34^+^ cells). The following mouse monoclonal antibodies were added to the cells and incubated on ice for 30 min: Anti-CD3 PE-Cy5 (clone HIT3a; 1:200 dilution), Anti-CD4 PE-Cy5 (clone RPA-T4; 1:200 dilution) Anti-CD8 PE-Cy5 (clone RPA-T8; 1:200 dilution), Anti-CD11b PE-Cy5 (clone ICRF44; 1:200 dilution), Anti-CD14 PE-Cy5 (clone TuK4; 1:200 dilution), Anti-CD19 PE-Cy5 (clone HIB19; 1:200 dilution), Anti-CD20 PE-Cy5 (clone 2H7; 1:200 dilution), Anti-CD56 PE-Cy5 (clone B159; 1:200 dilution), Anti-CD235a PE-Cy5 (clone GA-R2(HIR2); 1:200 dilution), Anti-CD34 APC-Cy7 (clone 581; 1:25 dilution), Anti-CD38 APC (clone HIT2; 1:10 dilution), Anti-CD45RA BV785 (clone HI100; 1:25 dilution), Anti-CD90 FITC ( clone 5E10; 1:50 dilution), and Anti-CD123 PE (clone 7G3; 1:10 dilution). After the completion of the incubation, cells were washed with FACS buffer and pelleted by centrifugation (1,300 rpm x 5 min, 4°C). Stained cells were resuspended in FACS buffer and propidium iodide added immediately prior to FACS and manufacturer’s concentration.

*FACS analysis – ESAM expression validation*

Filtered and pelleted marrow cells were resuspended in ACK lysis buffer (1 mL) and allowed to incubate for 5 min at ambient temperature. Lysis was then quenched with 10 mL of FACS buffer and cells pelleted (1,300 rpm x 5 min, 4°C). RBC-depleted bone marrow cell pellets were resuspended in 800 μL ice-cold FACS buffer per mouse. Miltenyi Lineage depletion beads were added (100 μL per mouse) and incubated for 10 min at 4°C. Upon incubation completion cells were loaded on to a Miltenyi MACS magnetic separation column and the column washed 3 x 3 mL FACS buffer. Flow-through was pelleted (1,300 rpm x 5 min, 4°C). Depleted cells were resuspended in FACS buffer (100 μL per mouse) and Anti-CD34 FITC (clone RAM34; 5.0 μg/mL final concentration) was added. Cells were incubated for 45 min on ice prior to addition of the remaining antibodies: Anti-Lineage Cocktail A700 (3 μL per mouse), Anti-cKIT APC-eFluor780 (clone 2B8; 2 μg/mL final concentration), Anti-Sca1 PE-Cy7 (clone D7; 2 μg/mL final concentration), Anti-CD150 BV421 (clone TC15-12F12.2; 2 μg/mL final concentration), Anti-Flt3 PerCP-eFluor710 (clone A2F10; 2 μg/mL final concentration), Anti-IL7Ra BV711 (clone SB/199); 2 μg/mL final concentration), Anti-CD16/32 PE (clone 93; 2 μg/mL final concentration), and Anti-ESAM APC (clone 1G8; 2μg/mL final concentration). The cells were incubated for an additional 30 min on ice with the complete cocktail prior to washing with FACS buffer to remove excess antibody. Cell were pelleted (1,300 rpm x 5 min, 4°C) and resuspended in fresh FACS buffer containing DAPI (1:1000 stock solution) prior to FACS.

*Fluorescence microscopy*

Cells were purified as described above and spun in a Cytospin centrifuge onto superfrost plus glass slides (10 min x 1000 rpm). Upon spin completion, slides were dried (5 min) and a circle drawn around the cells with a wax pen. Fixation buffer (4% PFA in PBS) was added on top of the cells and allowed to incubate (10 min). Fixative was pipetted away and the cells washed with PBS (3 x 5 min). Cells were incubated with permeabilization buffer (0.1% Triton X-100 in PBS, 10 min) after which time permeabilization buffer was removed and replaced with blocking buffer (5% donkey serum in 0.1% TritonX-100 in PBS). Cells were incubated in blocking buffer for 16 h at 4 ℃. Primary antibody (Pfkl) was added (1:100) in blocking buffer and incubated for 2 h. Upon completion slides were washed with PBS (3 x 5 min) and appropriate secondary added (1:500 in blocking buffer). Secondary was allowed to incubate for 1 h prior to PBS washes (3 x 5 min). DAPI (1:100, final concentration 200 ng/mL in PBS) was added to the slides and allowed to incubate for 10 min prior to mounting.

*Processing of purified cell types for mass spectrometry analysis*

Sorted cells were pelleted and washed 2x with ice-cold PBS to remove any remaining FBS (1,300 rpm x 5 min 4°C). PBS was aspirated away, and the pellets were snap frozen with liquid nitrogen prior to storage at -80°C. Prior to lysis, cells were thawed on ice and subjected to sample preparation with the PreOmics iST NHS kit according to literature protocol. The only modification made to the protocol was scaling down in volume of lysis buffer and digest buffer (20 μL each). Samples were resuspended in 12 μL of LC-Load Buffer from the iST NHS kit and peptide concentration determined (Pierce Quantitative Colorimetric Peptide Assay).

*Mass spectrometry analysis – liquid chromatography and timsTOF Pro*

A nanoElute was attached in line to a timsTOF Pro equipped with a CaptiveSpray Source (Bruker). Chromatography was conducted at 40°C through a 25cm reversed-phase Aurora Series C18 column (IonOpticks) at a constant flow-rate of 0.4 μL/min. Mobile phase A was 98/2/0.1% Water/MeCN/Formic Acid (v/v/v) and phase B was MeCN with 0.1% Formic Acid (v/v). During a 120 min method, peptides were separated by a 4-step linear gradient (0% to 15% B over 60 min, 15% to 23% B over 30 min, 23% to 35% B over 10 min, 35% to 80% over 10 min) followed by a 10 min isocratic flush at 80% for 10 min before washing and a return to low organic conditions. Experiments were run as data-dependent acquisitions with ion mobility activated in PASEF mode. MS and MS/MS spectra were collected with *m*/*z* X00 to 1500 and ions with *z* = +1 were excluded.

*Mass spectrometry data analysis*

Raw data files were processed with Byonic software. Fixed modifications included +113.084 C. Variable modifications included Acetyl +42.010565 N-term, pyro-Glu -17.026549 N-term Q, pyro-Glu -18.010565 N-term E. Precursor tolerance 30.0 ppm.

*Data compilation*

We generated lists of all UniProtIDs, gene names and their respective mappings that appear in all raw mouse cell files. All ‘nan’, ‘’ (empty strings), and ‘2 SV’ were ignored. We used the Retrieve/ID mapping program available at uniprot.org to find all possible gene name mappings for each UniProtID and all possible UniProtID mappings for each gene name. Using the data from the raw files and mappings from UniProt, we recursively mapped between UniProtIDs and gene names, 1) identifying all sets of UniProtID and gene name aliases, and 2) creating a list of unique mappings between every UniProtID aliases and gene name aliases. Any unnamed set of aliases were given a protein ID (UNM #) or a gene name (Unm #). For each given mapping between UniProtID aliases and gene name aliases, a single UniProtID alias and gene name alias were selected. During compilation of data for analysis, each UniProtID and gene name from a raw file was replaced by the selected aliases for consistency.

*RNA Isolation and Library Preparation*

RNA was isolated with TRIzol as per the manufacturer’s recommendations and was further facilitated by using linear polyacrylamide as a carrier during the procedure. We treated the total RNA samples with RQ1 RNase free DNase to remove minute quantities of genomic DNA if present. DNase treated samples were cleaned up using RNAeasy minelute columns. 1-10 ng of total RNA was used as input for cDNA preparation and amplification using Ovation RNA-Seq System V2. Amplified cDNA was sheared using Covaris S2 using the following settings: duty cycle 10%, intensity 5, cycle/burst 100, total time 5 min. The sheared cDNA was cleaned up using Agencourt Ampure XP. 500 ng of sheared cDNA were used as input for library preparation using NEBNext Ultra DNA Library Prep Kit for Illumina as per manufacturer’s recommendations.

*RNA-Seq and Data Analysis*

Libraries were sequenced using NextSeq 500 (Illumina) to obtain 2x150 base pair paired-end reads and HiSeq 2000 to get 2x100 base pair paired-end reads.

*Data normalization*

All RNA-sequencing data was normalized to transcripts per million (TPM). All individual mass spectrometry runs were normalized to sum to 1,000,000 (referred to as the “processed data”) to mirror the standard TPM used in RNA-sequencing analyses. For combined analyses (ex. protein vs mRNA), cells of the same type were averaged across all runs of the processed data (referred to as the “combined processed data”).

*PCA*

For PCA, the processed data was normalized such that the sum of values across cells for each gene was equal for all genes. PCA was performed on this data using the pca package available in scikit-learn. The 1-dimensional plot of each component direction was normalized by subtracting the entire data by the minimum value and dividing by the new maximum value. The list of genes and their relative contribution to each component is available in Table 4.

*Comparison between protein and mRNA*

The non-linear relationship between protein and mRNA when visualized on a plot prompted us to use spearman correlation to determine the monotonic relationship between protein and mRNA. As we were interested in the overarching view of protein regulation differences between cells, we only included genes expressed in both the proteome and transcriptome of all cell types. We then normalized these genes to sum to 1,000,000 per cell type before performing the spearman correlation test using spearmanr package available in scipy.

*GMM*

Protein comparisons

For each cell type from the combined processed data, the log_2_ fold change was determined against all other cell types for genes that were commonly expressed within every comparison. The values were combined into a single vector and used to run the GaussianMixture package available in scikit-learn. The model was fit 1,000 independent times. The two gaussians from each fit was grouped into either a low-variance gaussian or a high-variance gaussian. Each fit consistently yielded one low-variance and one high-variance gaussian. To determine the final two gaussians, the parameters (mean, variance and weight) in each group were averaged across all gaussians in the group.

Protein/mRNA comparisons

For each cell type from the combined processed data, the log_2_ fold change was determined against their respective bulk mRNA data for genes that were expressed both as protein and mRNA. The difference was taken for this value against those of all other cell types, and the differences were combined into a single vector. All subsequent steps were identical to that of protein comparisons following the establishment of a single vector for a given cell type.

**QUANTIFICATION AND STATISTICAL ANALYSIS**

Cell-to-cell analyses were performed on GraphPad Prism. One- and two-way ANOVA, unpaired and multiple t-tests were used where appropriate. Global analyses of proteomic and bulk RNA-seq data were performed via Python 2.7.15. Numpy 1.15.3 was used for vector operations, and matplotlib 2.2.3 was used for the generation of graphs. Statistical tests (spearman & pearson correlation), dimensionality reduction and distribution modeling (GMM) were carried out using scipy 1.2.1 and scikit-learn 0.20.3. Center was measured using the mean. Dispersion was measured using SD (standard deviation). Where appropriate, 99.7% confidence interval (3 SD) or 95% confidence interval (2 SD) was used with justification.

Use of 1.5 fold change as a threshold for identifying proteins of interest yielded too many proteins to be able to feasibly study for our initial analysis. We established a systematic, interpretable method of identifying proteins of interest. Use of QQ-plots and their pearson correlations (using pearsonr package available in scipy) when compared to a null distribution (generated from a Gaussian with equal center and dispersion as the data) consistently informed us that the log2 fold change data was not normal, and that it was better fit with a mixture of two gaussians—one with low variance and another with high variance. We assumed the low-variance gaussian (with higher weight) to be the large set of genes non-differentially expressed, but with technical noise. We assumed the high-variance gaussian (with lower weight) to be the smaller subset with more biological relevance. Our threshold for identifying proteins of interest was determined either by 3 SD away from the mean of the low-variance Gaussian (to exclude proteins identified due to noise), or by 2 SD away from the mean of the high-variance Gaussian (to determine biological significance)—whichever was more extreme. We validated our analysis at several steps using the housekeeping gene, Hprt1.

**DATA AND CODE AVAILABILITY**

The datasets generated during this study are available as supplemental tables. Code used for data compilation and analyses, as well as all other generated files not explicitly used or mentioned in this study, is available at https://github.com/jnoh4/PofHemat.
