## Supplemental Figures for "Mass spectrometry analysis of mouse hematopoietic stem cells and their progenitors reveals differential expression within and between proteome and transcriptome throughout adult and aged hematopoiesis"

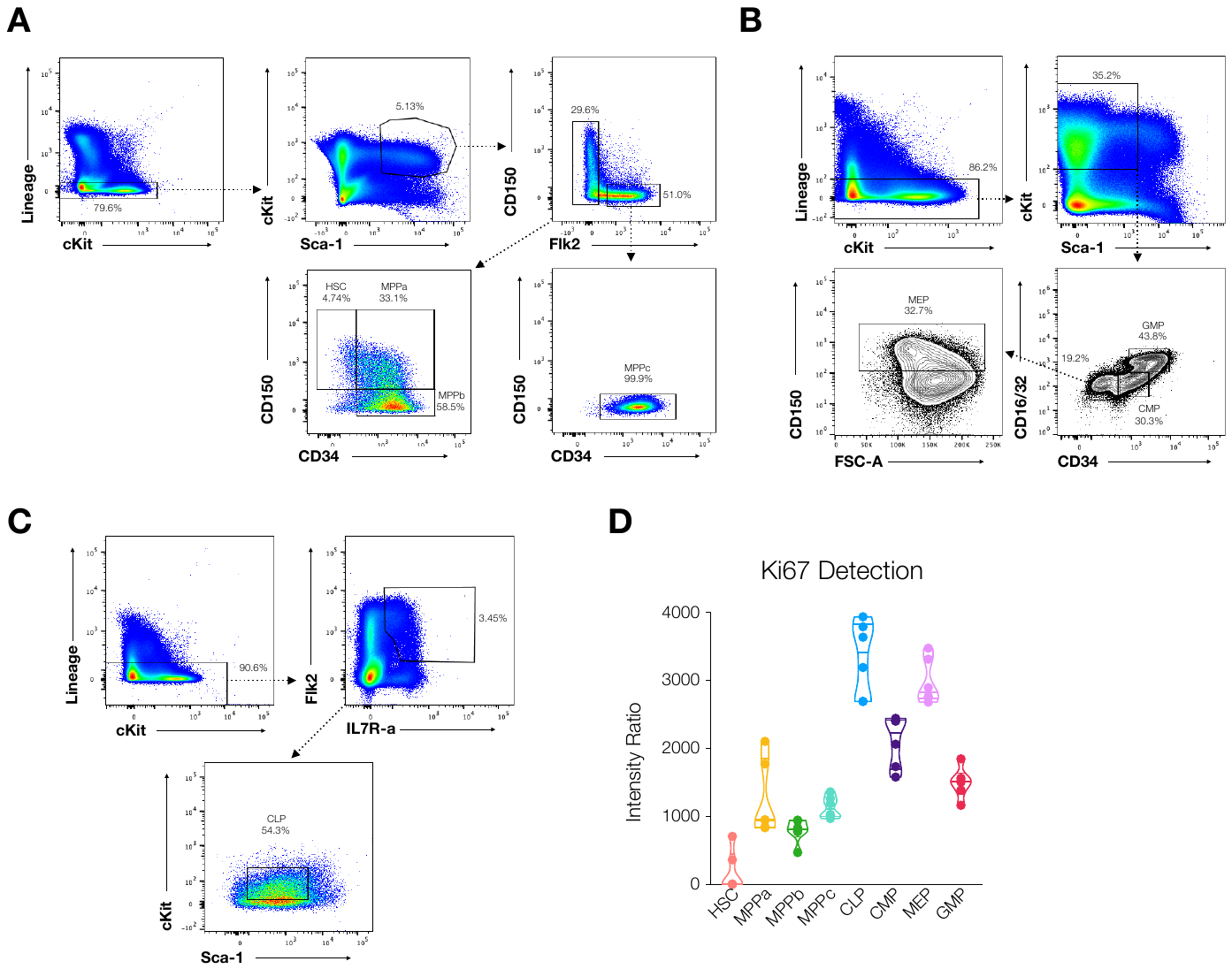

Supplemental Figure 1. Validation of pure cell populations. A. Representative sorting scheme for HSCs and MPPs. HSC (Lin^-^, cKIT^+^, Sca1^+^, CD34^-^, CD150^+^, Flt3^-^) MPPa (Lin^-^, cKIT^+^, Sca1^+^, CD34^+^, CD150^+^, Flt3^-^) MPPb (Lin^-^, cKIT^+^, Sca1^+^, CD34^+^, CD150^-^, Flt3^-^) MPPc (Lin^-^, cKIT^+^, Sca1^+^, CD34^+^, CD150^-^, Flt3^-^). B. Representative sorting scheme for GMPs (Lin^-^, cKIT^+^, Sca1^lo/-^, CD34^hi^, CD16/32^hi^), CMPs (Lin^-^, cKIT^+^, Sca1^lo/-^, CD34^med/hi^, CD16/32^-/lo^) and MEPs (Lin^-^, cKIT^+^, Sca1^lo/-^, CD34^-^, CD16/32^-/lo^, CD150^+^). C. Representative sorting scheme for CLPs (Lin^-^, CD34^med/hi^, Flt3^+^, IL7Rα^+^, cKIT^lo^, Sca1^lo^). D. Violin plots for intensity ratios of Ki67 across 6 replicates per cell type.

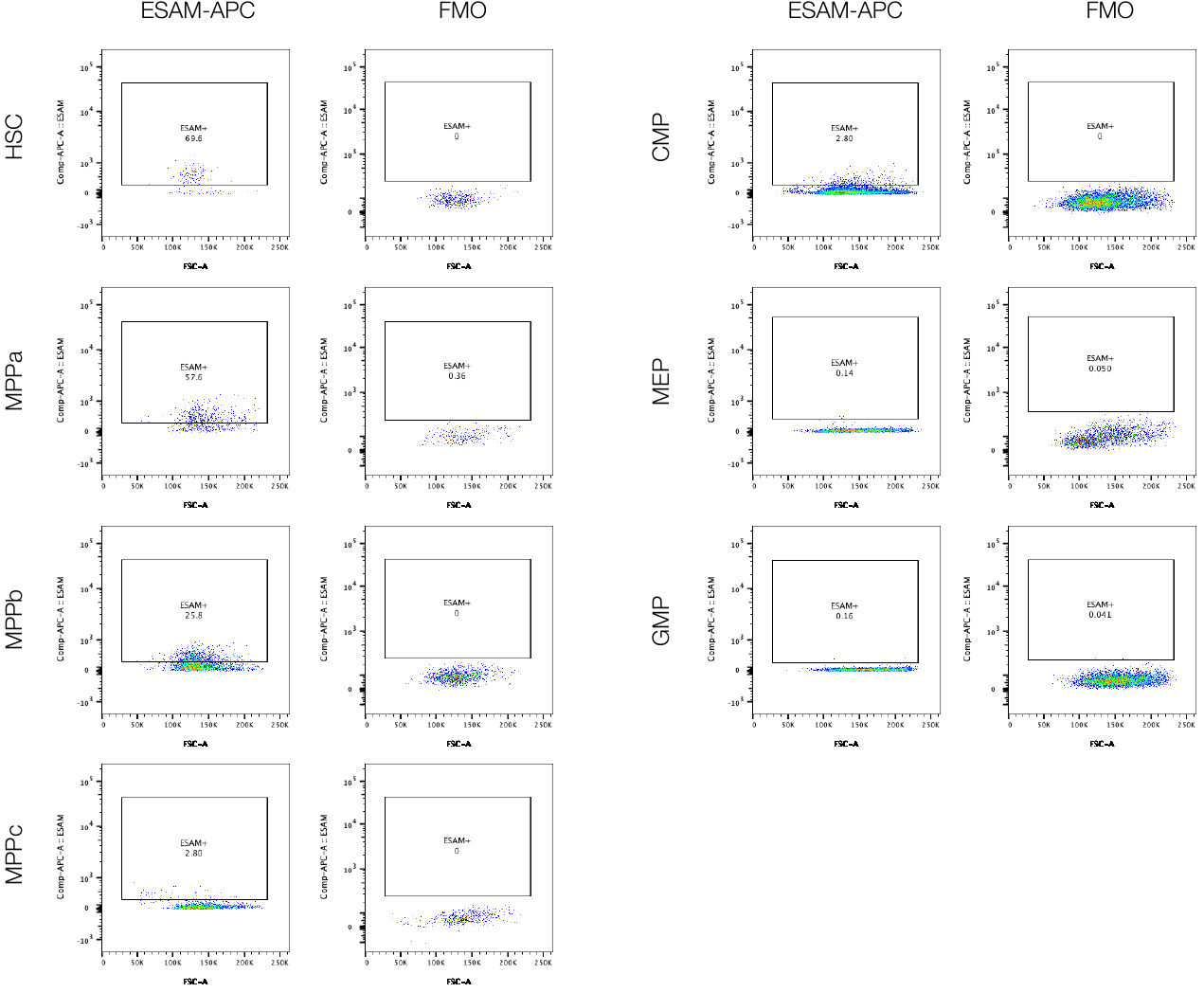

Supplemental Figure 2. Representative gating scheme for ESAM expression. N = 5 mice.

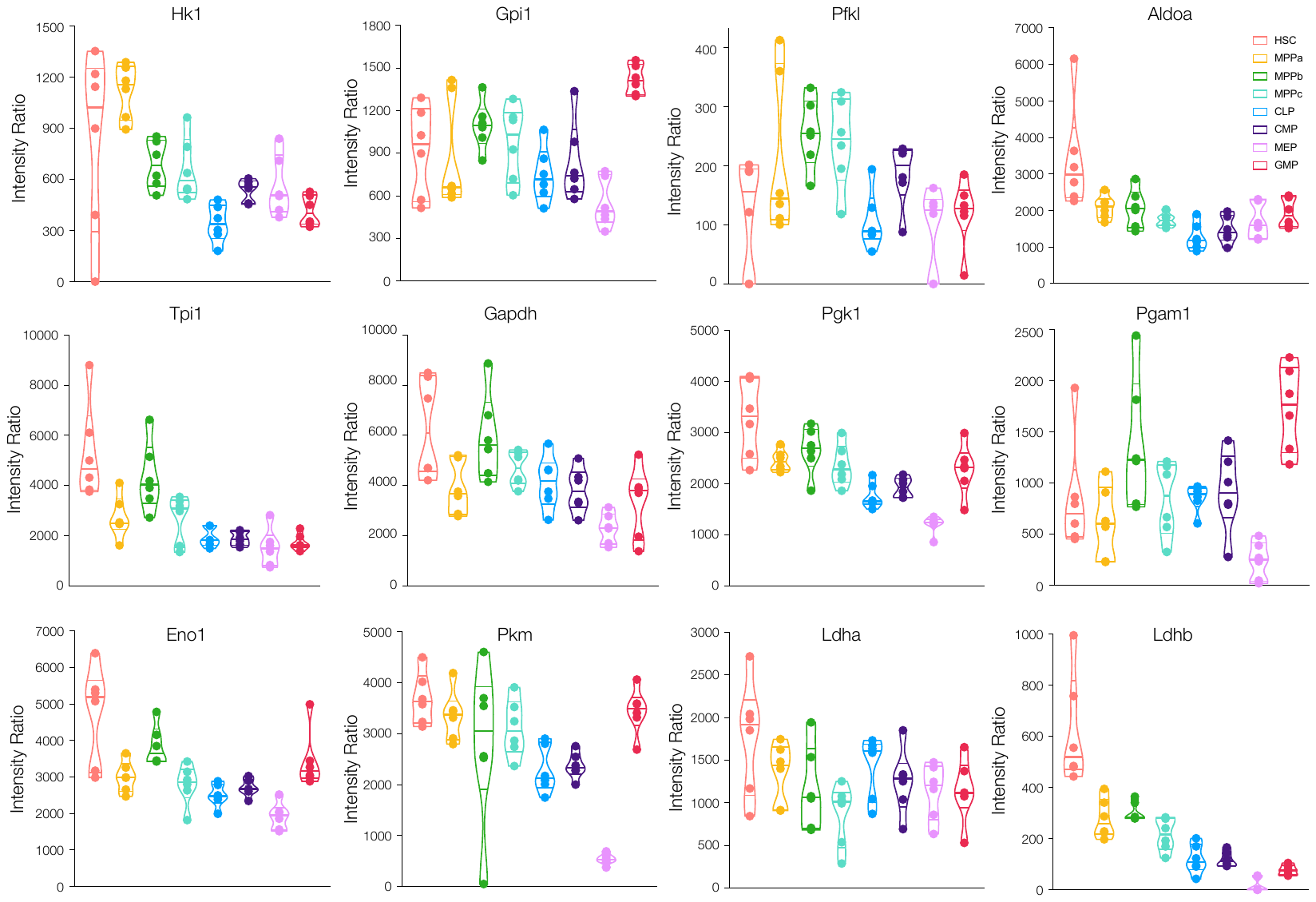

Supplemental Figure 3. Protein expression of glycolytic enzymes in adult HSCs and progenitors. Violin plots for intensity ratios of Hk1, Gpi1, Pfkl, Aldoa, Tpi1, Gapdh, Pgk1, Pgam1, Eno1, Pkm, Ldha, and Ldhb across 6 replicates per cell type.

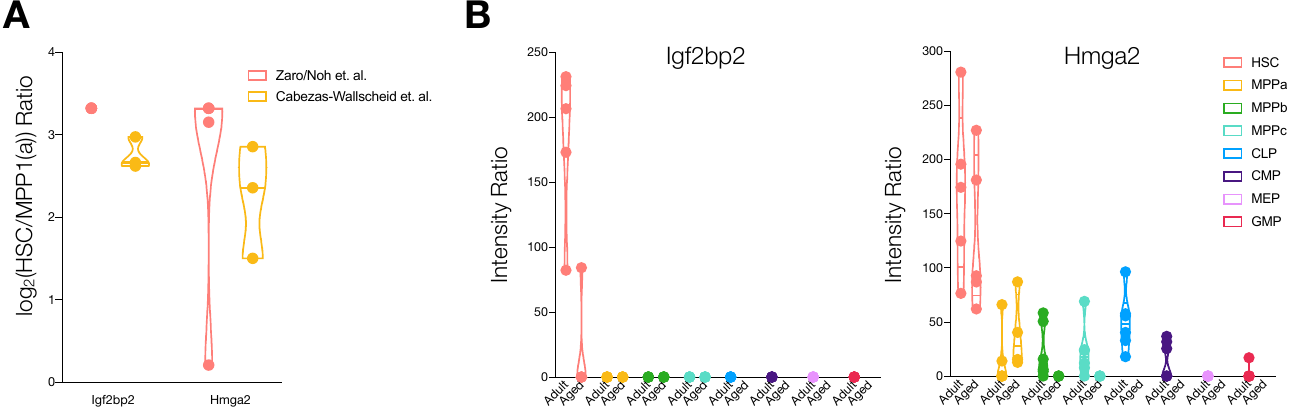

Supplemental Figure 4. Comparison of proteomic data from Cabezas-Wallscheid et. al. A. Enrichment ratios between HSCs vs. MPP1 or MPPa (log_2_) for Igf2bp2 and Hmga2. N = 3. Ratio maximums was set at 10 (log_2_(10) = 3.32. B. Protein expression of Igf2bp2 and Hmga2 in adult HSCs and progenitors. Violin plots for intensity ratios across 6 replicates per cell type.

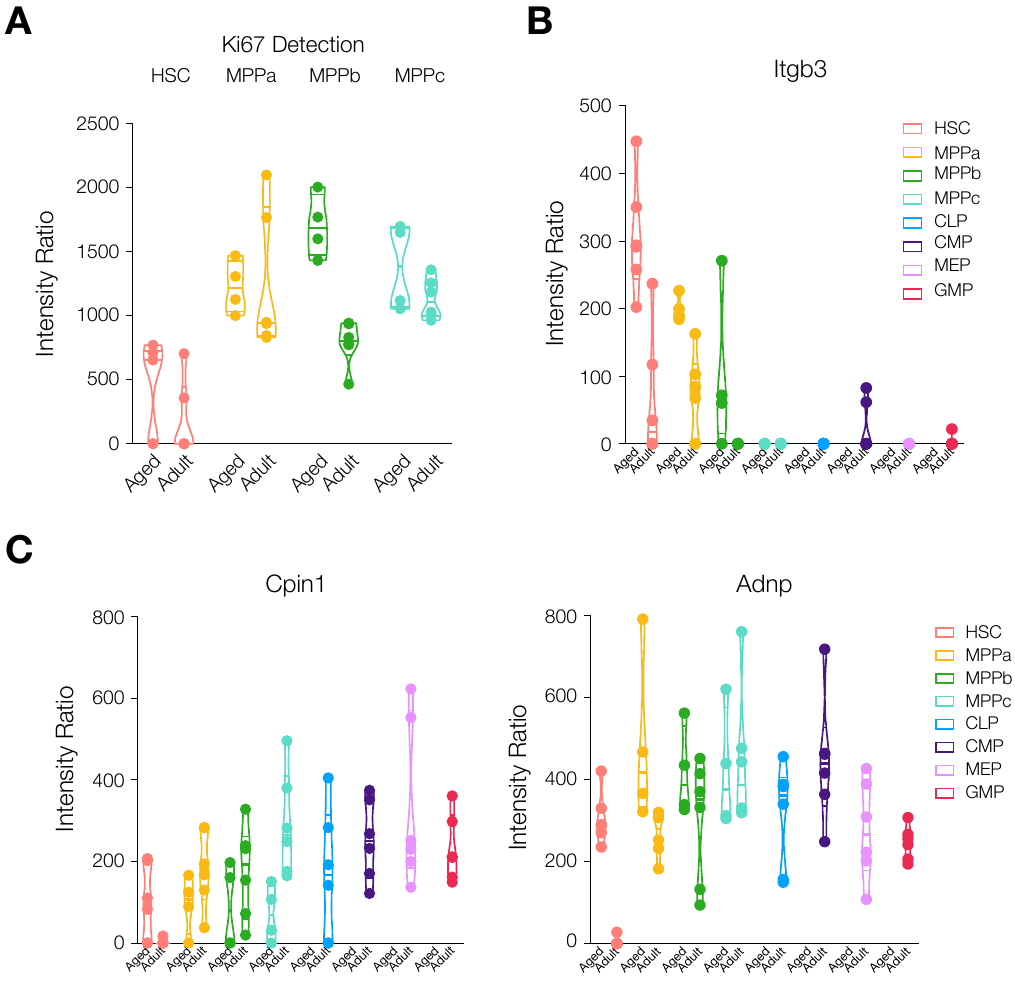

Supplemental Figure 5. Protein expression in aged HSCs and progenitors. A. Protein expression of Ki67 in adult and aged HSCs and progenitors. N = 6 for adult cells and aged HSCs. N = 4 for aged MPPs. B. Protein expression of the age-associated protein Itgb3 in adult and aged HSCs and progenitors. N = 6 for adult cells and aged HSCs. N = 4 for aged MPPs. C. Protein expression profiles for Cpin1 and Adnp. Violin plots for intensity ratios of Cpin1 and Adnp across all cell types. N = 6 for adult cells and aged HSCs. N = 4 for aged MPPs.

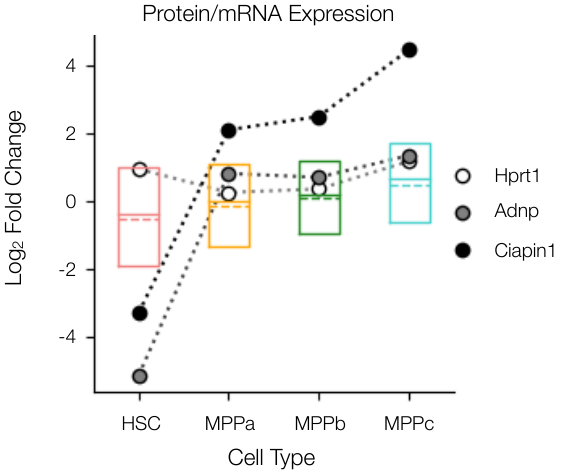

Supplemental Figure 6. Log_2_ fold change between protein vs. mRNA expression for proteins expressed across all 4 cell types and specific proteins of interest Adnp and Ciapin1 and the housekeeping gene Hprt1.
